# Disentangling sRNA-Seq data to study RNA communication between species

**DOI:** 10.1101/508937

**Authors:** JR Bermúdez-Barrientos, O Ramírez-Sánchez, FWN Chow, AH Buck, C Abreu-Goodger

## Abstract

Many organisms exchange small RNAs during their interactions, and these RNAs can target or bolster defense strategies in host-pathogen systems. Current sRNA-Seq technology can determine the small RNAs present in any symbiotic system, but there are very few bioinformatic tools available to interpret the results. We show that one of the biggest challenges comes from sequences that map equally well to the genomes of both interacting organisms. This arises due to the small size of the sRNA compared to large genomes, and because many of the produced sRNAs come from genomic regions that encode highly conserved miRNAs, rRNAs or tRNAs. Here we present strategies to disentangle sRNA-Seq data from samples of communicating organisms, developed using diverse plant and animal species that are known to exchange RNA with their parasites. We show that sequence assembly, both *de novo* and genome-guided, can be used for sRNA-Seq data, greatly reducing the ambiguity of mapping reads. Even confidently mapped sequences can be misleading, so we further demonstrate the use of differential expression strategies to determine the true parasitic sRNAs within host cells. Finally, we validate our methods on new experiments designed to probe the nature of the extracellular vesicle sRNAs from the parasitic nematode *H. bakeri* that get into mouse epithelial cells.

## INTRODUCTION

Organisms do not live in isolation. The wonderful diversity and complexity in life arises in part due to the contacts that living beings have with their peers. Symbioses can have positive or negative consequences to one or both of the interacting partners. These interactions are not only obvious at the macroscopic level, but molecular exchanges underlie many of them. Molecules moving between organisms of different species include antibiotics, toxins, volatiles, sugars, amino acids, amongst many others.

RNA is a molecule of incredible functional versatility, participating in central cellular processes as messenger, transfer and ribosomal RNA, but also in complex regulatory layers, from bacterial riboswitches to eukaryotic microRNAs (miRNAs). Yet RNA has historically been regarded as an unsuitable molecule for exchanging signals between cells or organisms due to its instability, even though it was proposed as an extracellular communicator several decades ago [1].

Recent advances in sequencing technology have allowed researchers to measure RNAs with unprecedented sensitivity, leading to the surprising discovery that many small RNAs, including miRNAs, are extracellular components of many human bodily fluids like blood, tears and maternal milk [2–4]. These extracellular RNAs can be protected from degradation through binding to proteins like Argonaute and/or encapsulation within extracellular vesicles (EVs) [5]. Even so, a report that miRNAs from plant food sources could be detected in the mammalian bloodstream was quite surprising [6]. These so-called “xenomiRs” have been hotly debated, with a slight consensus arising that miRNAs detected after passing through the vertebrate digestive tract are probably contaminations or other molecular errors coming from index swapping during Illumina library preparation [7–9].

Interestingly, a key discovery came when *Botrytis cinerea*, a fungal plant pathogen, was shown to secrete small RNAs, that traffic into plant cells to help block the host defense response [10]. Since then, we and others have shown that small RNAs are detected in material exchanged between a large variety of pathogens and their hosts [11–15]. The parasitic nematode *Heligmosomoides bakeri* secretes small RNAs inside EVs into the host gut environment, modulating the immune response of mice [11]. The parasitic plant *Cuscuta campestris* produces miRNAs that travel into the host tissue eliciting a functional silencing response in *Arabidopsis* [13]. Plants can also deliver their own sRNAs to strike back at their pathogens [14,15]. RNA exchange even occurs between the different domains of life: the bacterium *Salmonella* uses the host Argonaute to produce miRNA-like RNA fragments that increase its survival [16], and mammalian miRNAs present in the gut can be internalized into bacteria and affect their growth thereby shaping the microbiota [17]. Although there has been more focus in the literature on RNA released from pathogens, RNAs are probably being exchanged between all sorts of other interacting species [17–20].

Sequencing technologies are at a state were detecting RNAs of different sizes, from all sorts of biological material, even single cells, is accessible to most research groups. Analysis of RNA sequencing of interacting organisms began a few years ago, with “Dual RNA-Seq” experiments that focused on transcriptional analyses of bacterial pathogen-host systems [21,22]. To successfully perform these experiments, several technical aspects needed to be addressed to account for highly abundant rRNA or tRNA from phylogenetically heterogeneous samples, the lack of poly-A tails in prokaryotes, and scenarios where one of the organisms is present in very small relative amounts. In contrast, the bioinformatic analyses of these results are almost straightforward, since 100-150 nt sequences (the most common read-length of current Illumina sequencers) can usually be easily assigned to the correct position, in the correct genome of origin.

Dealing with eukaryotic small RNAs (∼20-30 nt) presents completely different challenges. Removal of rRNA and tRNA, or poly-A selection is not required, since a simple size-selection step prior to, or after, library generation will enrich for the RNA population of interest. On the other hand, bioinformatic analyses can be challenging since very short sequences can map perfectly to a large genome just by chance. Furthermore, short sequences can map to multiple locations, leading to uncertainty that is sometimes solved by discarding these sequences. Some sequences can also genuinely arise from different species. Ancient miRNAs, as well as highly conserved rRNA/tRNA fragments can share up to 100% identity between phylogenetically diverse organisms like nematodes and mammals. On the other hand, new miRNAs are constantly evolving, and they have been proposed as phylogenetic markers [23]. Taking advantage of this idea, miRTrace was developed to predict the taxonomic diversity in any sRNA-Seq sample or detect the origin of cross-species contaminations [24]. Yet because it relies on a database of clade-specific miRNAs, it cannot classify sequences that have not been curated.

There is increasing interest in studying the small RNAs that are naturally exchanged between organisms. Recently we discovered that the extracellular vesicles (EVs) secreted by the parasitic nematode *Heligmosomoides bakeri* are enriched in 5’ triphosphate sRNAs derived from repetitive elements [25], and not mostly microRNAs as we had found initially using standard library preparation techniques [11]. This is quite significant, since the sRNAs secreted by *Botrytis cinerea* that impair plant defense responses derive from transposable elements [10]. It is possible that many pathogens use repetitive elements of their genome to efficiently explore a wide range of sequences to interfere with their hosts. There are no available methods to confidently detect and quantify these kinds of sRNAs within the cells or tissues of another organism. Here we describe the development of methods to detect, quantify, and characterise sRNAs that can move between different species.

We downloaded available data from experiments designed to probe inter-organismal communication mediated by small RNAs. To further increase our dataset diversity and address scenarios where there are very low levels of parasite sRNAs, we designed new experiments to discover which of the sRNAs in *H. bakeri* EVs actually get into mouse host cells. We detail the difficulties of analysing these kinds of experiments, and propose a series of strategies to solve them. One of the biggest challenges arises from the sRNAs which can confidently map to the genomes of both interacting species. We show that this ambiguity depends on the length of the sRNA, the size of the genomes, and their phylogenetic relationship. We next demonstrate how sequence assembly of the raw sRNA-Seq data extends the length of many sRNAs and reduces the ambiguity problem. Finally, we show how differential expression analysis, in combination with sRNA assembly, and proper experimental designs, can be leveraged to confidently detect and quantify the sRNAs that move between even closely related species.

## RESULTS AND DISCUSSION

### A diverse selection of species that exchange small RNAs

To build a foundation for bioinformatically probing cross-species RNA communication, we selected sRNA-Seq samples representing interactions from four phylogenetically diverse pairs of organisms (**Table 1**). These were the model plant *Arabidopsis thaliana* infected by a fungus (*Botrytis cinerea*) or a parasitic plant (*Cuscuta campestris*), and the mongolian gerbil (*Meriones unguiculatus*) infected by a filarial parasite (*Litomosoides sigmodontis*). Given our interest in parasitic nematodes and their secreted extracellular vesicles (EVs), we also designed new experiments to explore the EV small RNAs from *Heligmosomoides bakeri* that get internalized by host cells, using sRNA-Seq of a mouse intestinal epithelial cell line. The full list of sRNA-Seq samples available from these experiments are described in **Supplementary Table 1**.

**Table 1.** Small RNA sequencing datasets of interacting organisms

| Host | Parasite | Tissue or condition | Data availability | Reference |
| --- | --- | --- | --- | --- |
| *Arabidopsis thaliana* | *Botrytis cinerea* | Rosette leaves: 24, 48 and 72 hours after infection | Short Read Archive: SRP019801. Samples: SRX252403, SRX252404, SRX252405 | [10] |
| *Arabidopsis thaliana* | *Cuscuta campestris* | *Arabidopsis* stems with an attached *Cuscuta* haustorium | Short Read Archive: SRP118832. Samples: SRX3214810, SRX3214811 | [13] |
| *Meriones unguiculatus* | *Litomosoides sigmodontis* | Serum from infected gerbils | GEO: GSE112949. Samples: GSM3091975, GSM3091976, GSM3091977, GSM3091978, GSM3091979 | Quintana, *et al.* 2019 |
| *Mus musculus* | *Heligmosomoides bakeri* | MODE-K cell line: 4 and 24 hours after adding EVs | GEO: GSE124506. Samples: GSM3535462, GSM3535463, GSM3535464, GSM3535468, GSM3535469, GSM3535470 | This work |

Since the field of cross-species communication by RNA is still young, these represent some of the only real-world scenarios of symbiotic models that have been examined with sRNA-Seq. The biological material sampled in each case is diverse: infected stems or leaves in the case of *Arabidopsis*, serum from infected gerbils, and a cell culture for our nematode-mouse model. The amount of parasite RNA present within the infected samples can also be quite different. *Botrytis* spores are used to infect *Arabidopsis* leaves, from which RNA is extracted after the necrotrophic fungus has grown and invaded the tissue of its host. For *Cuscuta,* we selected the samples that included the haustorium connected to *Arabidopsis* stems. As such, both *Arabidopsis* experiments included parasite tissue, and not only extracellular material. For the rodent models, on the other hand, the pathogen releases extracellular RNA to the host environment and the parasites are not themselves present in the collected material. We expect these last samples in particular to be akin to a “needle in a haystack” problem, with very small amounts of parasite sRNA amongst a very large amount of host RNA. In contrast, the plant samples are expected to contain a mixed population of parasite and host sRNAs at more comparable levels.

### Determining the amount of host, parasite, and ambiguous reads in sRNA datasets

As a first step to identify the genome of origin of sRNAs involved in cross-species communication, we prepared a combined reference genome for each pair of interacting species (see Methods, and **Supplementary Table 2**). We then focused only on sRNA reads between 18-30 nucleotides that map with 100% identity to the corresponding combined reference. These mapped reads are then divided into three categories: i) host (if they only map to the host portion of the reference), ii) parasite (if they only map to the parasite) and iii) ambiguous (if they map at least once to the host and at least once to the parasite). With this partitioning, different experiments yield varying proportions of host, parasite and ambiguous reads (**Figure 1**).

**Figure 1.**
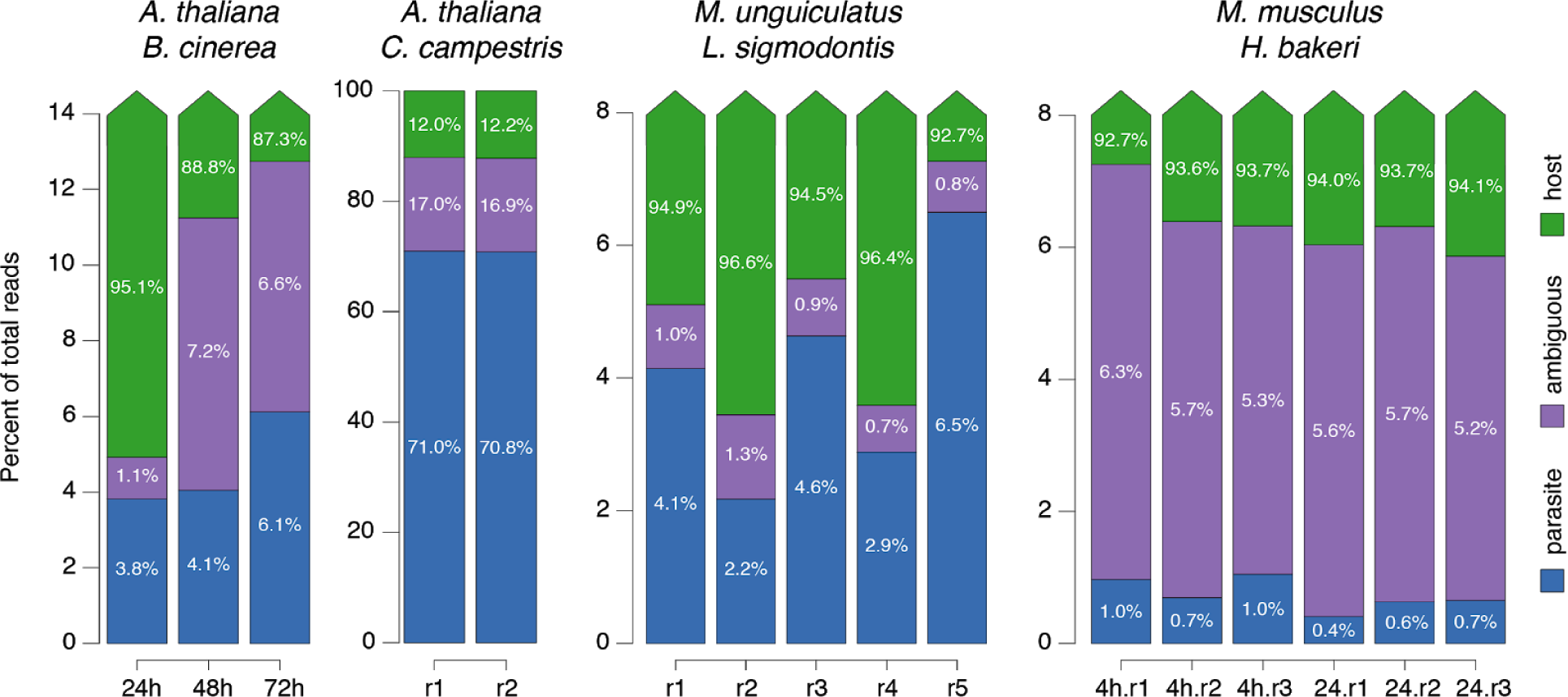
Percent of ambiguous and parasitic reads from infected samples. Each column represents one sRNA-Seq sample, and columns are grouped by experiment. Each experiment is given the name of the two interacting species, and the Y-axis is scaled independently to highlight the percent of parasite (blue) and ambiguous (purple) reads. Host reads (green) in all cases make up the remainder of 100%. Biological replicates are defined by “r” and time points post infection (*B. cinerea*) or incubation with EVs (*H. bakeri*) noted (further detailed in Table 1).

The *Arabidopsis + Botrytis* libraries show between 3.8-6.1% of parasite reads (increasing with the time post infection), with ambiguous reads accounting for 1.1-7.2%. The *Arabidopsis* + *Cuscuta* libraries show 71% of parasitic and 17% of ambiguous reads. The infected gerbil serum had between 2.2-6.5% of parasite reads but only around 1% of ambiguous reads. Finally, the MODE-K cells treated with extracellular vesicles from *H. bakeri* yielded the lowest amount of parasite reads, in the range of 0.4-1%. In this case, the parasitic reads are clearly outnumbered by the ambiguous ones, with 5.2-6.3% being assigned to this category. These results highlight the difficulty in correctly identifying all the parasitic sRNAs. Whilst one approach would be to discard the ambiguous reads we would in all cases be throwing away an important amount of sequencing information that may include bonafide RNA molecules involved in cross-species communication.

### Ambiguity in host-parasite sRNA-Seq reads is influenced by read length, genome size and phylogenetic distance

We next wanted to determine what factors lead to the ambiguous reads. There are at least three variables that could contribute: the length of the read, the size of the genomes, and the phylogenetic relationship of the genomes. We present these factors from a theoretical standpoint, using “k-mers” (nucleotide words of length *k*) as a proxy for reads.

#### Read length

Intuitively, it is more likely that a small k-mer will be present in two genomes compared to a longer k-mer. To illustrate this, we define two random genomes of the same sizes as *A. thaliana* and *B. cinerea*, and calculated the fraction of shared k-mers of different sizes [26]. The shared k-mers between these two random genomes decrease rapidly as *k* increases (**Figure 2a**). For instance, almost 80% of k-mers of length 12 are shared, but when considering k-mers of length 18, less than 0.003% are shared.

**Figure 2.**
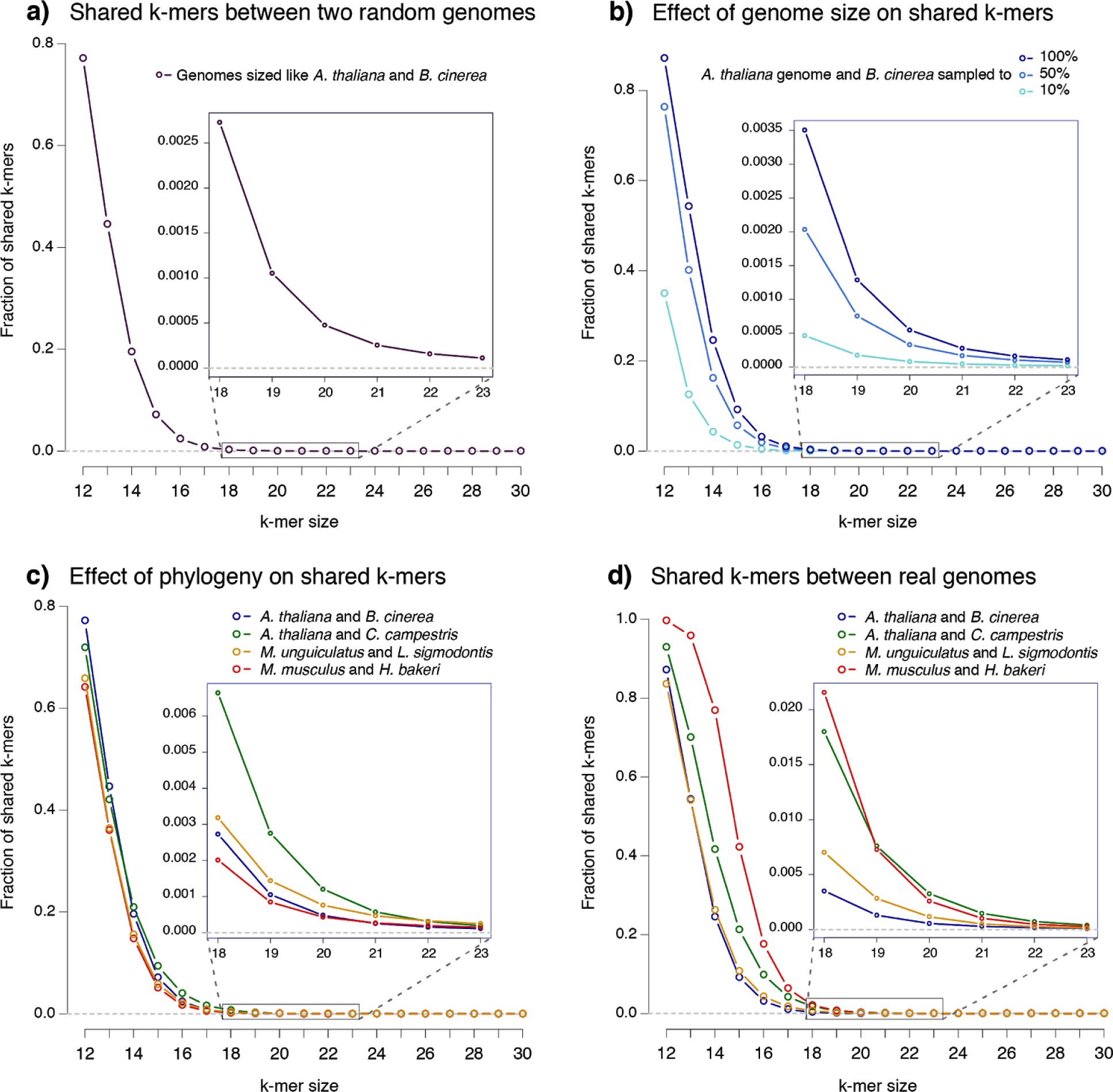
Factors that influence the number of ambiguous k-mers between pairs of genomes. X-axes represent the k-mer size and Y-axes the fraction of shared or ambiguous k-mers. a) Random genomes of sizes equivalent to those of *A. thaliana* and *B. cinerea*. b) Fixed *thaliana* genome, compared to full *B. cinerea* genome or a sample corresponding to 50% or 10% of the complete genome. c) All genomes are subsampled to the size of the smallest, that of *cinerea*. d) Real fractions of ambiguous k-mers in each pair of complete genomes. Insets correspond to a zoomed in area of k-mer sizes 18-23.

#### Genome size

The size of each genome determines the maximum number of distinct k-mers that it contains (**Supplementary Figure 1**). A smaller genome will have fewer distinct k-mers, and so the number of shared k-mers it can have with another genome is also expected to be smaller. To highlight this property, we took the real *A. thaliana* genome, but sampled decreasing fractions of the *B. cinerea* genome (100%, 50% and 10%) to visualize how the number of shared k-mers changes. As expected, smaller *Botrytis* genomes share a smaller percent of k-mers of any length (**Figure 2b**).

#### Phylogenetic distance

Real genomes are not random concatenations of nucleotides, but are related through shared ancestry. Thus, the phylogenetic distance between two genomes should also influence the number of shared k-mers and therefore our ability to distinguish small RNAs that might map to both. If we imagine two genomes that have just begun to diverge, almost all k-mers will be shared. To quantify the effect of phylogenetic separation, but ignoring the effect of genome size which we described above, we fixed the smallest of the genomes under consideration (*B. cinerea*) and randomly down-sampled each of the other six genomes to this size.

The effect of phylogenetic distance is small but noticeable (**Figure 2c**). In particular *A. thaliana* shares more k-mers with another plant (*C. campestris*) than with a fungus (*B. cinerea*). While both pairs of animal genomes are expected to be similarly related (rodents and nematodes), *H. bakeri* shares fewer k-mers with mouse than *L. sigmodontis* with the gerbil. This can be explained since *H. bakeri* has a particularly large genome (∼700Mb, compared to ∼65Mb for *L. sigmodontis*), that is full of repetitive elements many of which are unique to this species [25]. A random sample of the *H. bakeri* genome will thus include more k-mers from these repetitive elements. This helps explain the smaller fraction of shared k-mers than expected due to phylogeny, and highlights an extra contributing factor: genome composition and complexity, which we will not explore further in this work.

It is thus not possible to predict the exact number of ambiguous k-mers between two species just based on their genome size, but if the genomes are available it can be efficiently calculated using tools like Jellyfish [27]. By doing so, we can see that *H. bakeri* and *M. musculus* show the highest level of ambiguous k-mers, while *A. thaliana* and *B. cinerea* show the lowest (**Figure 2d**). These are the biggest and smallest pairs of genomes, respectively, indicating that genome size is a major factor driving these differences. But at longer k-mers the two plant genomes are the pair with the highest ambiguity. This is due to their close phylogeny (both species are eudicotyledons, a clade of flowering plants). In all cases the ambiguous k-mers between real genomes, at larger k-mer sizes, become much higher than expected exclusively by genome size, reflecting the contribution of shared ancestry (**Supplementary Figure 2**).

From these results there are two important things to note: 1) even for k-mers the size of biologically important molecules like microRNAs (∼21 nucleotides), there is always a fraction of sequence space that will be shared identically between two genomes, and 2) if we could increase the length of any sequence, even by one or two nucleotides, the probability that it will be shared between genomes would decrease substantially.

### High levels of ambiguity in host-parasite sRNA-Seq reads is caused by conserved sequences like ribosomal, transfer, and microRNAs

The levels of ambiguity in our real sRNA-Seq data are much higher than predicted by the fractions of k-mers shared between pairs of genomes. For instance, almost 17% of all 18-30nt reads from the *A. thaliana* and *C. campestris* interaction are ambiguous (**Figure 1**), while only 1.8% of k-mers of size 18 are shared between the genomes (**Figure 2d**). This implies that the sRNA-Seq reads are not produced randomly across the genome, but come from regions with a higher-than-average level of conservation. This is not surprising if conserved classes of RNA, like ribosomal RNA, are being sequenced. So, from what regions are the sRNA-Seq reads being produced, particularly the ambiguous ones? We sought to answer this, focusing on the *A. thaliana* and *C. campestris* interaction where the problem of ambiguous reads is most apparent (**Figure 1**).

We extracted all the ambiguous reads from libraries of *A. thaliana* stems with *C. campestris* primary haustoria attached (average of 917,669 from the two replicates) and tabulated them by length (**Figure 3a**). The length distribution is as expected for small RNAs enriched in Dicer products, with more reads between lengths of 21-24 nt. Plants usually show a peak of 21 nt enriched with miRNAs, and a peak of 24 nt enriched with siRNAs that target transposable elements. Interestingly, the ambiguous reads show a marked preference for 21 nt. We then traced where all the ambiguous reads mapped in *A. thaliana*, which is better annotated, and classified them according to the annotation of this genome (**Figure 3b**). Most of the ambiguous reads map to rRNA (25%) and miRNA (24%), with a small contribution of tRNA (0.7%). This is a clear enrichment above expected since rRNA, miRNA and tRNA together occupy less than 0.1% of the *Arabidopsis* genome, while comprising 58% of the ambiguous reads. Less than 1% map to other kinds of annotations, including introns, while the remaining 41% map to unannotated intergenic regions, which in total occupy 59% of the genome. Many plant miRNAs are highly conserved [28], so it is not surprising that most of the 21 nt ambiguous reads coincide with conserved and highly-expressed miRNAs like MIR159, MIR161 and MIR166. Ribosomal reads are more evenly distributed across all read lengths, suggesting that their presence is caused by low levels of random fragmentation that are unavoidable for such highly abundant molecules. Lastly, tRNAs are less evenly distributed, with a slight peak at 26 nt observable in this case, suggesting that specific fragments of tRNAs are being sequenced. Ribosomal and miRNA contribution is also the main explanation for the ambiguous reads in libraries collected from the *Cuscuta* stem above the primary haustoria, and from *Arabidopsis* stems sampled ∼4cm above the point of *C. campestris* haustorial attachment (**Supplementary Figure 3**).

**Figure 3.**
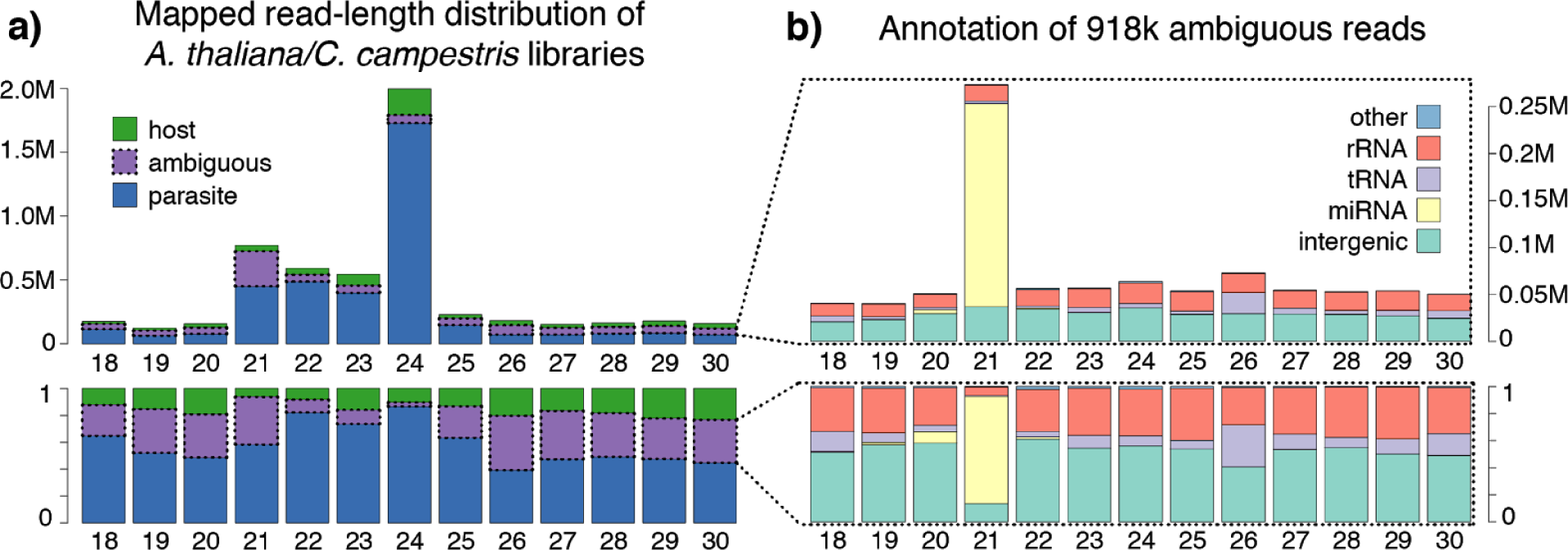
Genomic origin of ambiguous reads from libraries of *A. thaliana* stems with a *C. campestris* haustorium attached. Each bar represents the sequenced reads of one size between 18-30 nucleotides. Bar height represents the actual number of reads (top) or the fraction of reads (bottom). a) Mapping categories defined as in **Figure 1**, but split according to read length: host (green), parasite (blue) or ambiguous (purple). b) Genomic annotation of ambiguous reads only: rRNA (orange), miRNA (yellow), tRNA regions (light purple), intergenic (light green), or other annotation (light blue).

With these results we can see that discarding sRNA-Seq reads that map to rRNA and tRNA annotations, which is a common practice, can lead to a substantial reduction of ambiguity. Yet the ultimate goal of this work is to be able to detect RNA transfer between species and emerging literature suggests tRNA and rRNA fragments could be extracellular signaling molecules. For instance, tRNA fragments can be selectively packaged into extracellular vesicles and move between cells [29], while tRNA fragments in sperm can contribute to intergenerational inheritance [30]. So, there is likely to be important information that could be lost if we discard these kinds of sequences.

Discarding conserved miRNA sequences would be even more problematic, since foreign miRNAs are known to benefit from hijacking existing regulatory networks. A Kaposi’s sarcoma herpesvirus miRNA uses the same target site as the cellular miR-155 [31], while we have shown that nematode miR-100 and let-7, which are identical to their mouse counterparts, are present in secreted material during infection [11]. There is a need therefore to be able to track the origin of ambiguous sequences.

Even highly conserved miRNAs, tRNAs and rRNAs have point differences in some part of their sequence. For example, the loops of miRNA hairpins are poorly conserved. Due to the high depth of current sequencing technology, there will be reads with slightly different 5’ and 3’ ends, due to imperfect processing by Drosha and/or Dicer enzymes. There will also be reads overlapping the miRNA loop region. In fact, the existence of reads from different parts of the miRNA hairpins is the basis of popular prediction tools like miRDeep2 [32]. Thus, as long as we are able to extend the conserved sequences into a less conserved portion, we should be able to disambiguate them. We therefore explore the possibility of using sRNA-Seq assembly to reduce ambiguity through extension of reads.

### Assembly of sRNA-Seq reads

Most work on RNA sequence assembly has focused on producing full-length transcripts from mRNA-Seq data. There are many methods that work in a genome-guided fashion: first mapping reads to the genome, then assembling clusters (exons) and connecting them with rules based on splicing properties and sequencing depth, e.g. Cufflinks [33] and Stringtie [34]. Analogous to these, there are some tools that cluster sRNA-Seq reads where they map to the genome, in order to predict sRNA-producing loci: segmentSeq [35], CoLIde [36], and ShortStack [37]. ShortStack fits our needs quite well, since it analyzes reference-aligned sRNA-Seq reads to cluster them in order to predict sRNA genes, which we shall refer to as genome-guided clusters from here on. So, we used ShortStack to perform a genome-guided sRNA assembly and quantification.

We were also particularly interested in finding out if we could deal with situations in which the genomes for the interacting organisms were not available, or were not of sufficient quality. In these cases a *de novo* assembly approach is the only option. There has been a lot of development regarding *de novo* RNA-Seq assemblers. These tools do not require genome sequences, but rely instead on breaking down reads into k-mers, building a graph, and finding paths through the graph to build longer sequences. To our knowledge, these RNA-Seq *de novo* assemblers have not been used before on sRNA-Seq data. This makes sense, since k-mers of at least 25 nucleotides are usually used to improve the assembly quality, while functional molecules in sRNA-Seq (e.g. miRNAs) are generally smaller than this size. In our case, though, we want to extend sRNA sequences beyond the mature RNA, in order to capture sequence variation that can help us infer the correct genome of origin.

We tested six popular *de novo* transcriptome assemblers: Oases [38], rnaSPAdes [39], SOAPdenovo [40], Tadpole [41], TransABySS [42] and Trinity [43]. These programs first generate contigs by extending k-mers in a graph. This step produces short contigs that are later connected into full-length transcripts, but for our purpose of slightly extending sRNAs it could be sufficient, so we included the output of this “k-mer extension” step as a standalone method when possible (see Methods). One of the most important parameters for all the assemblers is the k-mer size, which affected the number of reads that we could remap to the assembly (**Supplementary Figure 4**). The optimal k-mer was 19 for all our sRNA-Seq datasets, except for the *A. thaliana* + *C. campestris* data where 21 was slightly better.

The four assemblies generated with only the first “k-mer extension” step (rnaSPAdes-only-assembler, Tadpole, Trans-ABySS-stage-contigs and Trinity-inchworm) performed quite differently than the full pipelines (**Supplementary Figure 5)**. They generated a larger number of contigs (**Figure 4a**), that were shorter (**Figure 4b**), and mapped better to the reference genomes (**Figure 4c**) than the full transcriptome assemblers. Additionally, library re-mapping was higher than with the other evaluated assemblies (**Figure 4d**). From these, Trinity-inchworm showed the highest library re-mapping across the evaluated datasets and is therefore used in our subsequent analyses.

**Figure 4.**
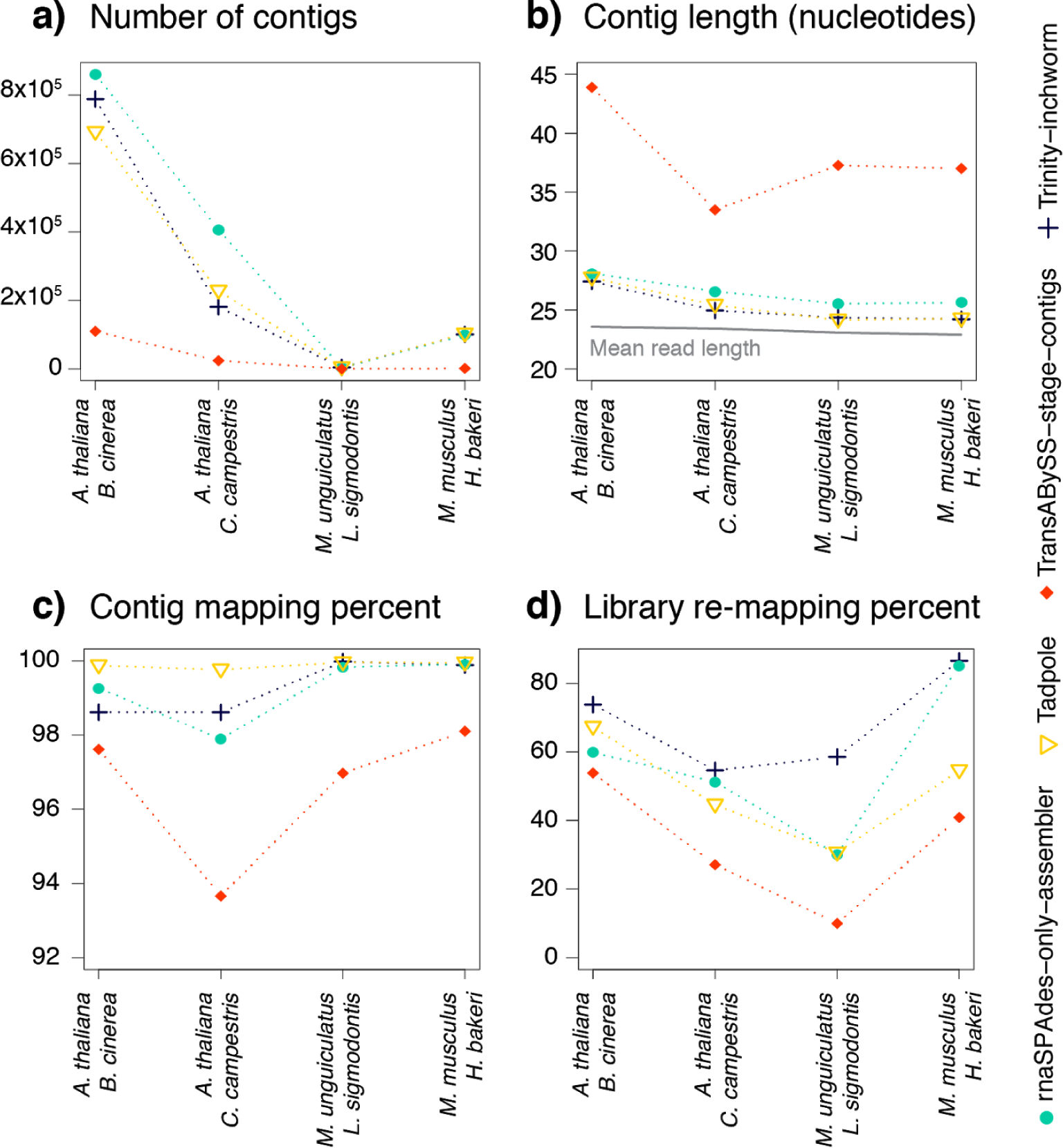
Evaluation of sRNA-Seq *de novo* assembly by “k-mer extension”. We compared the methods using the following criteria: a) total number of generated contigs, b) average length of contigs, c) percent of contigs that map perfectly to the reference genomes, and d) percent of initial libraries that map perfectly to the contigs. As a guideline, the mean library read length is shown in b). The full pipelines are compared in **Supplementary Figure 5**.

### Assembly reduces ambiguity of host-parasite sRNA-Seq reads

To compare the amount of ambiguity between the original reads (unassembled), *de novo* contigs and genome-guided clusters, we first assigned contigs (when possible) and clusters to their genome of origin. We then mapped reads directly to the sequences of the contigs or clusters. We used the number of uniquely-mapping reads in each contig or cluster to help distribute the reads that could map equally well to more than one contig or cluster (see Methods). For this analysis our only question was if the reads could be assigned confidently to one of the two interacting genomes. With either type of assembly, many reads previously annotated as ambiguous can now be assigned to one of the two interacting genomes (**Figure 5**). In general the contigs appear to be more conservative, with a modest increase in the parasite component, and maintaining a relatively large portion of ambiguous reads. The genome-guided clusters, on the other hand, include higher percentages of parasite reads than the *de novo* contigs which is in part due to the fact that some reads will be distributed randomly if no uniquely mapping reads are found nearby (detailed further below).

**Figure 5.**
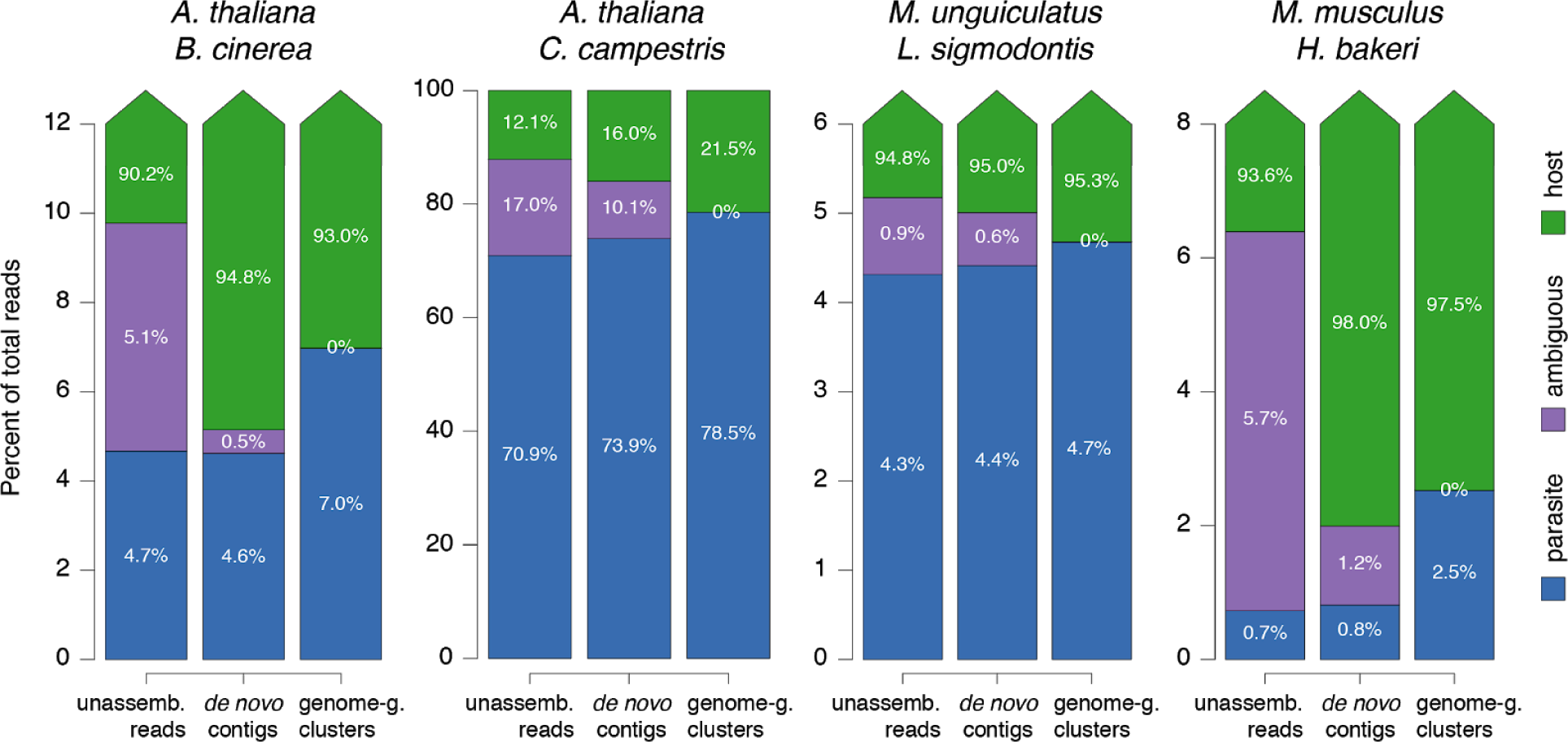
Percent of ambiguous and parasitic reads before and after assembly. Each column represents the average of sRNA-Seq samples for each experiment, first according to unassembled reads, then *de novo* contigs, then genome-guided clusters. Each experiment is given the name of the two interacting species, and the Y-axis is scaled independently to highlight the percent of parasite (blue) and ambiguous (purple) reads. Host reads (green) in all cases make up the remainder of 100%.

These results show how the assembled versions of the sRNA-Seq data contain more reads that can be assigned to the interacting organisms, and less ambiguity, allowing researchers to use more information from their experiments. The recommended assembly strategy (*de novo* or genome-guided) will depend on particular experiments. If one or both genomes are not available or are not of sufficient quality, *de novo* assembly is the best alternative. For high quality genomes, genome-guided assembly yields even less ambiguity. However, the assembled sequences could include reads from the wrong genome, due to errors during assembly, or because ShortStack randomly distributes a certain number of ambiguous reads between the genomes. For these reasons, we ideally want an independent test for validating the origin of the assembled sequences mapped to the parasite genome.

### Differential expression analysis improves detection of parasite sRNAs

Ideally, parasite sRNAs should be present in those samples that were infected with the parasite, and be absent (no reads) in uninfected samples. Unfortunately, this does not perfectly hold up, due to problems like index-swapping during library preparation [44,45]. Especially for situations when the parasite sRNAs can be present in very low numbers, a statistical framework is needed to determine which sRNAs are reliably present in the infected compared to the uninfected samples. For this, we can use differential expression analysis, which also helps to confirm if our assembled sequences behave like parasite or host sequences.

We designed our new *H. bakeri* extracellular-vesicle (EV) experiment to be amenable to differential expression analysis. We collected RNA from three biological replicates of MODE-K intestinal epithelial cell cultures treated with *H. bakeri* EVs, and the corresponding untreated controls. Since we do not know the dynamics of import, or the stability of foreign sRNA once inside the cells, we performed RNA extraction at 4 and 24 hours after treatment and following extensive washing of cells. We then mapped all the sRNA-Seq reads to our assembled contigs and clusters, quantified their expression using ShortStack, and also obtained the simple counts of each unique unassembled read for the baseline analysis. For these three types of count matrices, we performed the exact same steps of a differential expression analysis (see Methods). We also kept track of *H. bakeri* and *M. musculus* mapping status for reads, contigs and clusters and used this information when visualising our results (**Supplementary Figure 6**). This helps us determine which reads/contigs/clusters may actually come from the host genome, despite mapping perfectly and preferentially to the parasite genome.

The process of sequence assembly reduces ambiguity, but another advantage is that it reduces the number of statistical tests during differential expression analysis (there are fewer distinct contigs/clusters than unassembled reads), reducing a problem known in Statistics as multiple-testing. In addition, if the reads are grouped correctly into real biological entities with a consistent expression pattern, we should get higher counts, which can increase statistical power.

Although we conservatively performed the differential expression analysis starting with all unassembled reads, contigs or clusters, we focused only on the subset that should contain the real parasite sequences: those that were assigned to the *H. bakeri* genome (parasite), and that were up-regulated in the EV-treated samples. With these criteria, the parasite sequences we detected with each strategy, included an average of 11,508 counts for the unassembled reads, 23,553 counts for the *de novo* contigs, and 64,729 counts for the genome-guided clusters. These results show how the assemblies have increased the number of confidently detected parasitic sequences: the *de novo* contigs contain twice as many counts, and the genome-guided clusters about 5.6 times more counts, compared to the unassembled reads (**Supplementary Table 4**).

Our mapping results (**Figure 5**) had indicated that all sequences that map perfectly to the parasite genome represent genuine parasite sRNAs. Our differential expression results suggest that in all cases they can still be divided into those that are genuine parasite sRNAs (up-regulated in samples treated with parasite EVs), and those that more likely represent host sRNAs (similar expression levels in treated and control samples). Nevertheless, the differential expression analysis could be underpowered (due to a small number of replicates and high biological variability) leading to false negative predictions. So we next wanted to further validate these results.

### Validation of differentially expressed parasitic sRNAs

A distinctive property of *H. bakeri* EV sRNAs is that the majority are 22-23 nucleotides in length and begin with a Guanine [25]. This is in stark contrast to endogenous MODE-K sRNAs that are dominated by miRNAs of 22 nucleotides that begin with a Uracil (**Supplementary Figure 7**). We thus have a simple method to determine whether there is a signature in the reads associated with true parasite sRNAs: compare the first-nucleotide preference of our predictions. We first classified all sRNA reads according to starting nucleotide and length, defining three categories: 22G (enriched in parasite EVs), 22U (enriched in MODE-K) or other (see Methods). As a reference, libraries prepared from untreated MODE-K libraries contain 2% 22G and 34% 22U, while pure *H. bakeri* EVs contain 62% 22G and <1% 22U reads (**Figure 6a**). The assembled contigs and clusters that do not show evidence of differential expression (non-DE) have high fractions of 22U reads, similar to mouse MODE-K libraries (**Figure 6b**). This would suggest that some of the assembled sequences are actually chimeras, i.e. they have incorporated a large number of sequences that are really from the host. Unfortunately we cannot rule out that some of these contain true parasite miRNA sequences that are diluted by the host content and remain as false negatives of our differential expression analysis. Nevertheless, all of the sRNAs that are significantly upregulated (DE) after treatment with parasite EVs are enriched with 22Gs, consistent with them being true parasitic sRNAs (**Figure 6b**). The unassembled reads classified as non-DE also have the 22G pattern of true parasitic sequences, but they are relatively few in number (6,928). Our proposed strategies show that the assembled contigs and clusters allowed us to discover a larger number of true parasitic sequences (25,555 22G reads for *de novo* contigs and 33,289 22G reads for genome-guided clusters) than the baseline analysis with unassembled reads (22,131 22G reads). In general, our results show that considering mapping information alone can be misleading, and that a differential expression approach is useful to separate parasite from host sequences, particularly for the assembled contigs and clusters.

**Figure 6.**
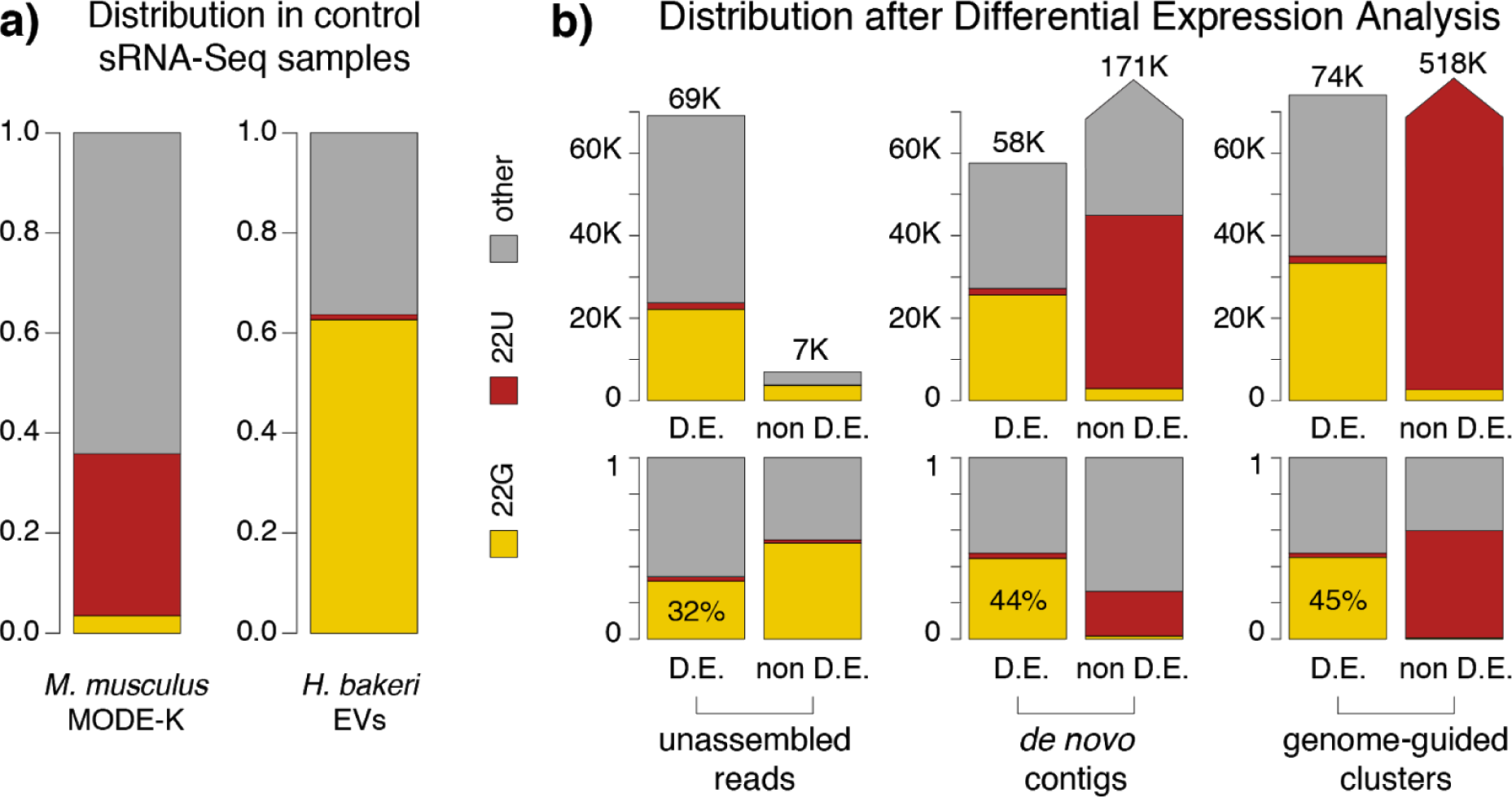
First-nucleotide categories for differential expression results. Reads were categorized as 22G (yellow), 22U (red), or other (grey) based on length and first-nucleotide. a) sRNA profiles of control samples: untreated MODE-K cells and purified *H. bakeri* Extracellular Vesicles (EVs). Bar height represents the fraction of all reads. b) sRNA profiles of unassembled reads, *de novo* contigs and genome-guided clusters. For each of these sets there are two bars, the first one represents differentially expressed up-regulated elements (D.E.) and the second, elements that lack evidence for differential expression (non D.E.). Bar height represents the number of reads (top) or the fraction of reads (bottom) belonging to these categories.

As a final validation of the parasite DE sequences that we detect inside host cells, we checked if they are also found in pure *H. bakeri* EV libraries. To do so, we first mapped all our pure *H. bakeri* EV reads to our DE contigs and clusters (for the unassembled reads, we checked which ones were identical). We do not expect to recover every sRNA read observed in EV libraries, since some EV sRNAs might not get into MODE-K cells, others might be turned over quickly or degraded, and others might not be detected due to insufficient sequencing depth. We reasoned, though, that the percent of recovered EV reads is an indication of how good the method is at recovering true parasite sRNAs within host cells. This analysis showed us that 1,811 DE unassembled reads correspond to 18.7% of the total reads in EV libraries, while 1,152 DE contigs and 1,432 DE clusters receive 29.9% and 42.3% of all EV reads, respectively (**Figure 7**). These results again highlight the improvement achieved by both assembly strategies, with the genome-guided clusters representing the best results according to all of our criteria.

**Figure 7.**
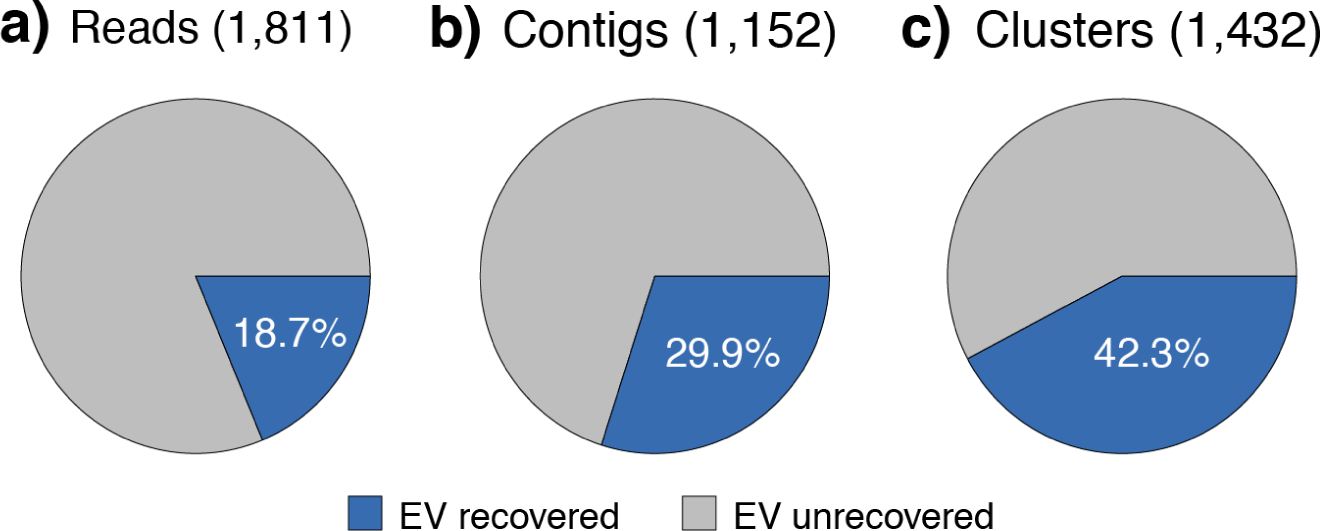
Percent of reads from pure *H. bakeri* EV libraries, recovered during differential expression analysis of MODE-K cells treated with EVs. Each circle represents the total reads from *H. bakeri* EV libraries (average of two replicates). The blue portions represent the fractions recovered by detected *H. bakeri* sequences according to each strategy, the grey portions represent EV reads that were not recovered. The number of *H. bakeri* differentially expressed elements detected with each strategy (up-regulated in cells treated with parasite EVs) is shown in parenthesis above each circle.

## CONCLUSIONS

We are now realising that the phenomenon of organisms exchanging RNA during their interactions is surprisingly widespread. These small RNAs can be produced and secreted by the cells of one organism, travel within extracellular vesicles, and perform regulatory functions when entering cells of a different species. We know very little about which kinds of RNAs can be secreted, which ones make it inside the cells of the receiving organism, and which have a functional role for the interacting organisms. We are just beginning to understand the potential functions and applications of this kind of RNA communication. Although the sequencing technology is at a state where we can begin to interrogate any pair of interacting species at unprecedented detail, there are no bioinformatic tools to correctly interpret the results. Before we can properly study the mechanisms and functions of RNA communication, we need to be able to correctly disentangle the sRNA-Seq data that is being acquired. We have shown here that the small size of sRNA-Seq sequences, and the large size of genomes, leads to many sequences mapping incorrectly or ambiguously to both interacting genomes (**Figure 2**). Even worse, many of the produced sRNAs that can be exchanged include sequences from highly conserved miRNAs, rRNA or tRNAs that are even more likely to map well to both genomes (**Figure 3**). We first showed that by performing sequence assembly of the sRNA-Seq data, we can greatly reduce the problem of ambiguity, and assign more sequences to their correct genome of origin (**Figure 5**). Importantly, we revealed that mapping information can still be misleading, and we showed that differential expression analysis can be used to confidently detect parasitic sRNAs that have been internalized by host cells.

We designed new experiments to detect the parasitic EV sRNAs from *H. bakeri* that successfully enter a mouse epithelial cell line. With our methods, we showed that 2% of the sRNA-Seq reads within treated MODE-K cells come from the parasite (**Supplementary Table 4**). This is a substantial increase over the simple approach of mapping to the genomes and dividing perfect hits between parasite and host, which suggested that only 0.7% of the sRNA-Seq reads were parasitic (**Figure 1**). Our genome-guided assembly increased this number to 2.5% (**Figure 5**) but we showed with differential expression that this was inflated with host sequences. The sequences within our final 2% estimate have all the characteristics of true *H. bakeri* EV sequences: they show the expected length and first-nucleotide 22G preference (**Figure 6**) and include more than twice the number of reads sequenced from independently purified EVs, compared to the approach using unassembled reads (**Figure 7**).

There are still some caveats to the methods we propose. Highly conserved sequences from the host, like miRNAs, can be misincorporated into parasitic sequence assemblies. The magnitude of this problem will depend on the relative level of expression of the conserved sRNA from both organisms in the sequenced sample. In our nematode-mouse experiment, a few miRNAs that we know are present in purified EVs (e.g. let-7, miR-100) are naturally expressed in MODE-K cells. Since even equally expressed sRNAs should be present in a ∼98/2 mouse/nematode ratio, it is not surprising that some mouse sequences erroneously contribute to the nematode assemblies. In any case, we believe that there is still room for improving sRNA-Seq assembly strategies. Promisingly, programs for *de novo* RNA-Seq assembly can be used, with appropriate parameters, and yield results that are comparable with genome-guided sRNA-Seq cluster assembly.

We have come to appreciate the great advantage of designing experiments to study RNA communication with differential expression in mind. Ideally this implies sampling from the separate organisms, and from the interacting material, all with several biological replicates. We realise that this might be a limitation in some cases, due to cost, the availability of sufficient quantity of biological material (e.g. purified EVs) or even the possibility of obtaining certain samples (e.g. from an obligate intracellular parasite). Nevertheless, we would like to stress the importance of having biological replicates and controls of at least one of the interacting organisms, particularly for confidently detecting low-abundance sRNAs.

Regardless of bioinformatic approaches, there may always be some sequences that are 100% identical between the interacting organisms. Careful use of chemically modified nucleotides might allow one to experimentally confirm the origin of some of these sequences. The most interesting next steps, though, will be to focus on understanding the function of the exchanged RNAs. Most work until now has focused on the assumption that extracellular sRNAs will behave as miRNAs when inside a different organism. Yet there are an increasing number of reports to suggest extracellular RNAs can operate through non-canonical mechanisms [46]. Furthermore, we have recently shown that *H. bakeri* EV sRNAs are mainly 5’ triphosphate species that are bound to a non-conventional worm Argonaute, which is unlikely to function like a miRNA Argonaute [25]. Here, we have now shown that these parasite sRNA sequences are stably detected inside mouse cells and future experiments will focus on understanding what these foreign RNA messages are doing to the host.

## METHODS

### Selected experiments and reference genomes

The list of host-parasite species used in this work is shown in **Table 1**. Further information of the sRNA-Seq data processed from these experiments is included in **Supplementary Table 1**. The reference genomes used are described in **Supplementary Table 2**. For each experiment, in addition to the separate reference genomes, a combined genome reference was produced, by concatenating the sequences from both genomes. In cases where rRNA was missing, these were manually added as an extra contig. A two word label was added to all fasta headers to differentiate parasite form host genome sequences. All genome files were indexed using Bowtie-1.2.2 [47].

### Small RNA-Seq library processing

The quality of all sRNA-Seq libraries was inspected using FastQC [48]. Raw reads were then cleaned and trimmed to remove 3’ adapter using reaper [49] with the following parameters: geom no-bc, mr-tabu 14/2/1, 3p-global 12/2/1, -3p-prefix 8/2/1, -3p-head-to-tail 1, -nnn-check 3/5, -polya 5 -qqq-check 35/10, -tri 35. Finally, only reads between 18-30 nucleotides were kept. When needed, reads were collapsed into individual sequences with counts, using tally [49]. One replicate of the MODE-K control cells (incubated for 24 hours without treatment) was an outlier according to PCA analysis, did not have a clear peak of mouse miRNAs (suggesting degraded RNA), and was excluded from further analyses.

### Calculations of host, parasite and ambiguous reads

Each library was aligned to the separate host and parasite genomes using Bowtie-1.2.2 [47] and requiring perfect end-to-end hits (-v 0). Each read was classified as: *host* if it only hit the host genome, *parasite* if it only hit the parasite reference and *ambiguous* if it hit both genomes.

### Shared k-mers between genomes

The fractions of k-mers between sizes 12-30 that are shared between each pair of genomes were calculated using Jellyfish 2.2.10 [27].

### Genome-guided sRNA assembly

To perform genome-guided sRNA assembly we used ShortStack 3.8.2 [37] with parameters favoring smaller clusters: a minimum coverage of one read, requiring 0 mismatches, using unique-mapping reads as guide to assign multi-mapping reads (mmap: u), a padding value of 1, reporting all bowtie alignments (bowtie_m: ‘all’), and a ranmax value of 5000 to avoid losing reads mapping to multiple sites. The default bowtie cores and sorting memory values were also increased to improve processing time. Reads were aligned to the concatenated host and parasite reference genomes described above.

### *De novo* assembly of sRNA-Seq

To evaluate *de novo* assembly of small RNA reads, six popular RNA-Seq *de novo* assemblers were selected: Oases [38], rnaSpades [39], SOAPdeNovo [40], Tadpole [41], TransAbyss [42] and Trinity [43]. These assemblers were also evaluated using only their first “k-mer extension” step: a) rnaSpades “--only-assembler”, Trans-AbySS “--stage contigs” and Trinity “--no_run_chrysalis”; b) the equivalent for Oases was to use contigs generated by velvetg, while for SOAPdenovo-Trans the. contig was used; c) Tadpole is a simple assembler that only performs k-mer extension. Additional parameters for each configuration are available in **Supplementary Table 3**. All the generated contigs were post-processed as follows: 1) all reads used to generate the assembly were aligned back to the contigs using Bowtie-1.2.2 (-v 0), and 2) using the BAM files from these alignments, contig edges that did not have any reads mapping to them were trimmed back. All contigs were then mapped to the reference genomes to decide if they were host, parasite or ambiguous, as described above.

### Disambiguation of host-parasite mixed samples

After applying the post-processing step to all contigs, reads that aligned to contigs were classified into three groups: reads that mapped to multiple contigs (*multi-mapping* reads), reads that mapped exclusively to one contig (*support* reads) and reads that did not align to any contig. In order to disambiguate some of the *multi-mapping* reads, the following criteria were used. Considering all contigs that received each *multi-mapping* read:

a. If the two contigs with the highest number of *support* reads were both from either *host* or *parasite*, the counts of the *multi-mapping* read were divided among the contigs according to the proportion of *support* reads.
b. If one of the two contigs with the highest number of *support* reads came from *host* and the other from *parasite*, the two numbers of *support* reads were compared to decide if they were significantly different, using a Poisson test (λ estimated as the average of these two numbers). The count of the *multi-mapping* read were distributed according to the proportion of *support* reads in each contig only if the difference of *support* read counts was significant (p-val < 0.05). If not, the counts were considered to remain ambiguous.

Additionally, all reads that did not align to any contig were collapsed to their unique sequences using tally. These new “unitigs” were concatenated to the contig file to be part of the reference. To define which sequences belonged to host, parasite or remain ambiguous, these sequences were aligned to both genomes using Bowtie-1.2.2. For a small number of contigs that did not align with Bowtie1 (-v 0), Bowtie2 was used with default parameters but allowing up to two mismatches (XM:i flag in the SAM file).

### *H. bakeri* life cycle and EV isolation

CBA x C57BL/6 F1 (CBF1) mice were infected with 400 L3 infective-stage *H. bakeri* larvae by gavage and adult nematodes were collected from the small intestine 14 days post infection. The nematodes were washed and maintained in serum-free media *in vitro* as described previously [25]. To collect *H. bakeri* EVs, culture media from the adult worms was collected from 24-92 hours post-harvest from the mouse (the first 24 hours of culture media was excluded due to potential host contaminants). Eggs were removed by spinning at 400 g and supernatant was then filtered through 0.22mm syringe filter (Millipore) followed by ultracentrifugation at 100,000 g for 2 h in polyallomer tubes at 4 °C in a SW40 rotor (Beckman Coulter). Pelleted material was washed two times in filtered PBS at 100,000 g for 2 h and re-suspended in PBS. The pelleted *H. bakeri* EVs, were quantified by Qubit Protein Assay Kit (Thermo Fisher), on a Qubit 3.0.

### MODE-K uptake assays

MODE-K cells were kindly provided by Dominique Kaiserlian (INSERM) and were maintained as previously described [50]. Uptake experiments were carried out with 2.5ug EVs per 50,000 cells for 4 and 24 hrs time points, in a 37 ^°^C, 5% CO_2_ incubator. Cells without incubating with *H. bakeri* EVs were treated as control with the two-time points. Cells were then washed twice in PBS before RNA extraction with a miRNeasy mini kit (Qiagen), according to manufacturer’s instructions. The RNA Integrity Number (RIN) was tested with the Agilent RNA 6000 Pico Kit on a Agilent 2100 Bioanalyzer. Three biological replicates were included for each of the samples.

### Small RNA sequencing

Total RNA was treated with RNA 5’ Polyphosphatase (Epicenter) following manufacturer’s instructions, before library preparation. Libraries for small RNA sequencing were prepared using the CleanTag small RNA library prep kit according to manufacturer’s instruction. For all samples, 1:2 dilutions of both adapters were used with 18 amplification cycles (TriLink BioTechnologies). Libraries of the length between 140-170bp were size-selected and sequenced on an Illumina HiSeq 2500 in high-output mode with v4 chemistry and 50bp SE reads, by Edinburgh Genomics at the University of Edinburgh (Edinburgh, UK).

### Differential expression analysis (DEA)

To perform differential expression analysis, a matrix was first built for individual sequences using all unique reads in the libraries to be compared. In this matrix rows represent individuals sequences and columns represent libraries. Each cell represents the times a sequence was found in a given library. A similar procedure was done to obtain matrices for contigs and clusters with the following modifications: each library was aligned to FASTA files of the contigs or clusters, and those reads that mapped to more than one sequence were distributed proportionally to *support* counts (see above).

Differential expression analyses were performed using the edgeR package [51]. The sRNA elements (individual sequences, *de novo* assembled contigs or genome-guided clusters) with low expression were filtered out: only those with more than one count per million in at least two libraries were kept. The MODE-K vesicle-treated libraries were compared to the control untreated MODE-K libraries, regardless of the incubation time (4 and 24 hours). To determine differential expression, a generalized linear model (GLM) likelihood ratio test was used, fixing a common dispersion value of 1.817 for reads, 2.141 for contigs, and 1.984 for clusters. False discovery rates (FDR) were calculated and sequences with a FDR lower or equal to 0.2 and a positive log fold-change were considered parasite sequences according to differential expression.

### Defining sRNA classes by length and first nucleotide

The first nucleotide and length of each sequence mapping to the *de novo* assembled contigs and genome-guided clusters was calculated using custom R scripts and the Rsamtools package. The criteria to classify a sequence as “22G” were: reads should begin with a Guanine, and be 22-24 nucleotides long. For the “22U” category: reads should begin with a Thymine and should be exactly 22 nucleotides long. These criteria were defined observing the properties of the pure EV and MODE-K libraries.

### Expression comparison with *H. bakeri* EV libraries

To quantify the expression in pure *H. bakeri* EVs of the contigs and clusters that were assembled using the infected MODE-K samples, the EV libraries were mapped onto the contigs and clusters using ShortStack. The same parameters were used as for defining sRNA clusters: minimum coverage of 1, perfect matches, unique-mapping reads as a guide to assign multi-mapping reads (mmap: u), a padding value of 1, and a ranmax value of 5000.

## ACKNOWLEDGEMENTS

We would like to thank Araceli Fernández Cortés for support using the Mazorka HPC cluster at Langebio. We also want to acknowledge Pablo Manuel González de la Rosa for help with running sRNA annotation pipelines and Sujai Kumar for suggesting and testing Tadpole for assembling sRNA-Seq data.

