## Supplementary Figures for "Disentangling sRNA-Seq data to study RNA communication between species"

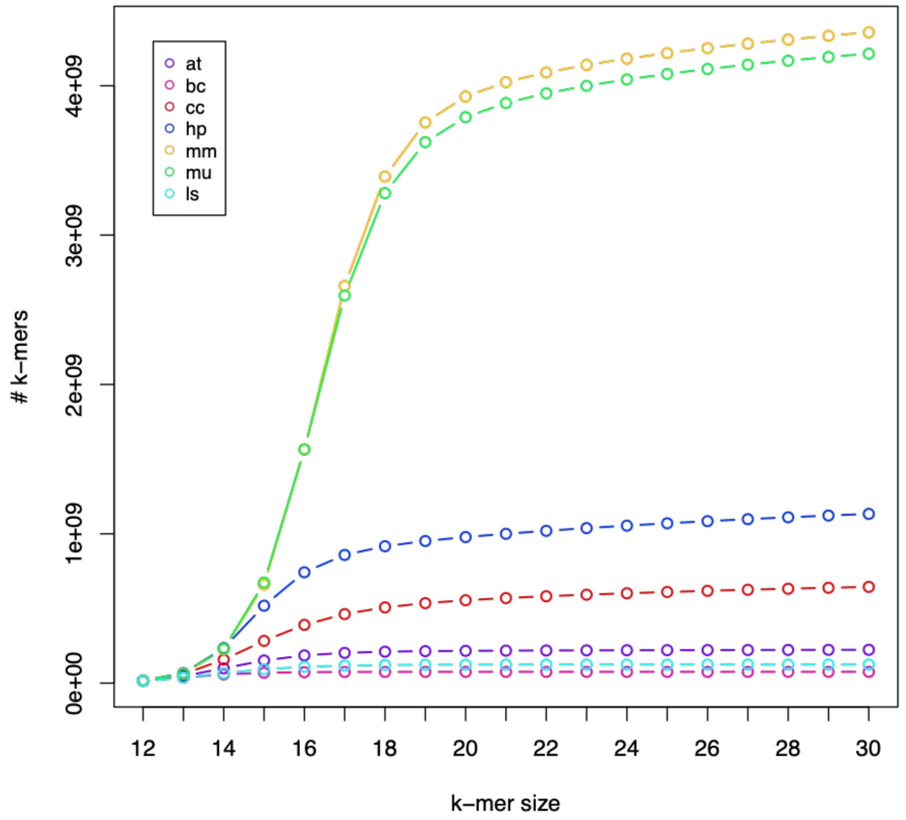


**Supplementary Figure 1. Number of k-mers in three host and four parasite genomes**. Each line represents a different species. at=*Arabidopsis thaliana*, bc=*Botrytis cinerea*, cc=*Cuscuta campestris*, hp=*Heligmosomoides bakeri*, mm=*Mus musculus*, mu=*Meriones unguiculatus*, ls=*Litomosoides sigmodontis*. Y-axis represents the total number of distinct k-mers of each size (X-axis) for each species.

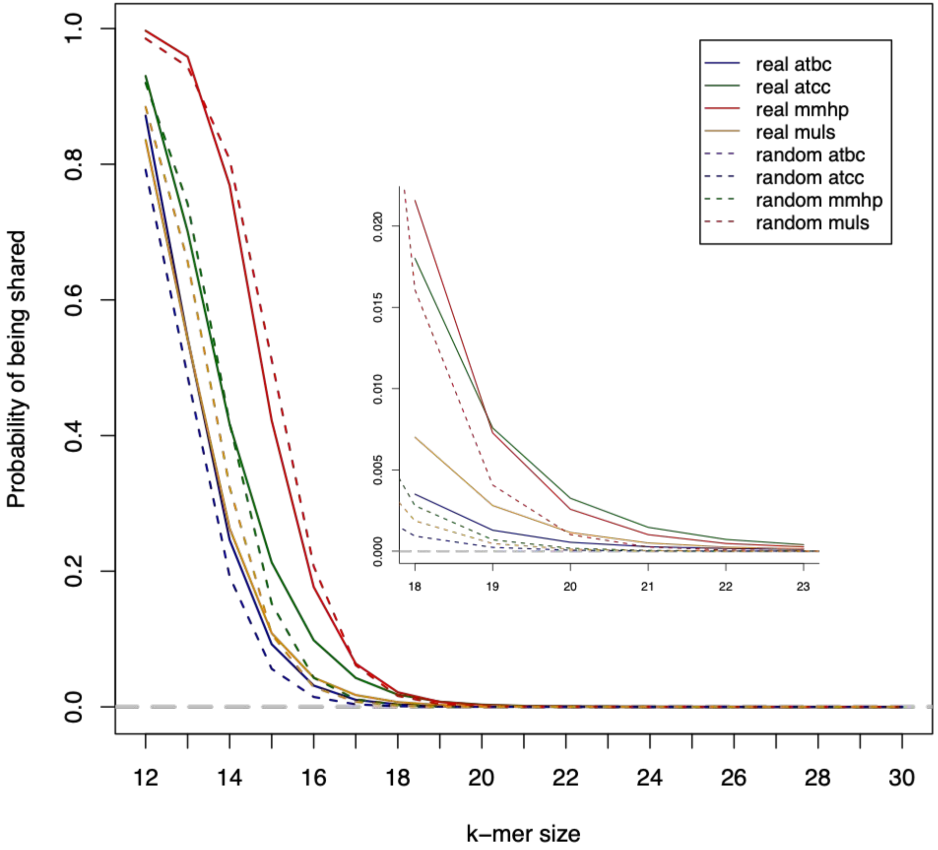


**Supplementary Figure 2. Fraction of shared (ambiguous) k-mers between pairs of interacting genomes.** Solid lines were calculated using real genomes whereas dotted lines were calculated for theoretical random genomes (see Methods). atbc=*A. thaliana* and *B. cinerea,* atcc=*A. thaliana* and *C. campestris*, mmhp=*M. musculus* and *H. bakeri*, muls=*M. unguiculatus* and *L. sigmodontis*. Inset correspond to a zoomed in area of k-mer sizes 18-23, where the real genomes clearly share more k-mers than the random versions.

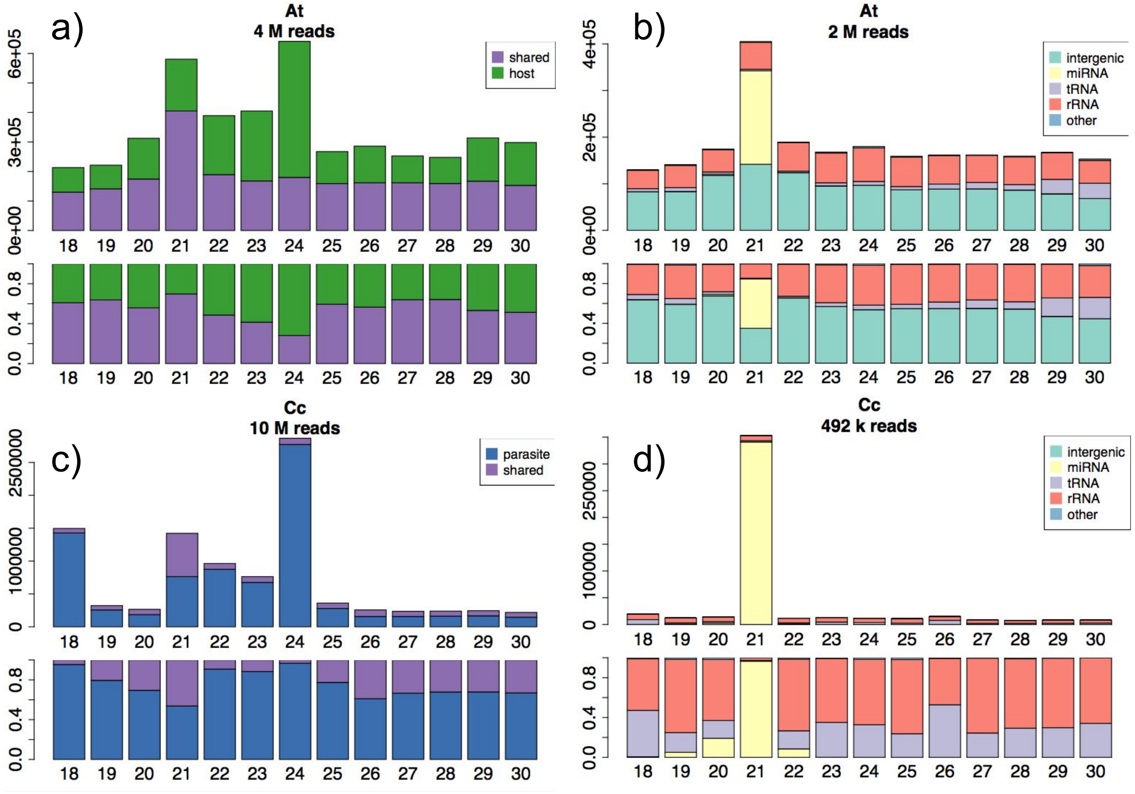


**Supplementary Figure 3. Genomic origin of ambiguous reads for *Arabidopsis thaliana* and *Cuscuta campestris* isolated libraries.** Each bar represents the sequenced reads of one size between 18-30 nucleotides. Bar height represents the actual number of reads (top) or the fraction of reads (bottom). a) and c) Read mapping categories split according to read length: host (green), parasite (blue) or ambiguous (purple). b) and d) Genomic annotation of ambiguous reads only: rRNA (orange), miRNA (yellow), tRNA regions (light purple), intergenic (light green), or other annotation (light blue). Libraries were made from a) and b) *Arabidopsis* stems ~4cm above the point of *C. campestris* haustorial attachment, and from c) and d) *Cuscuta* stems above the site of primary haustoria.

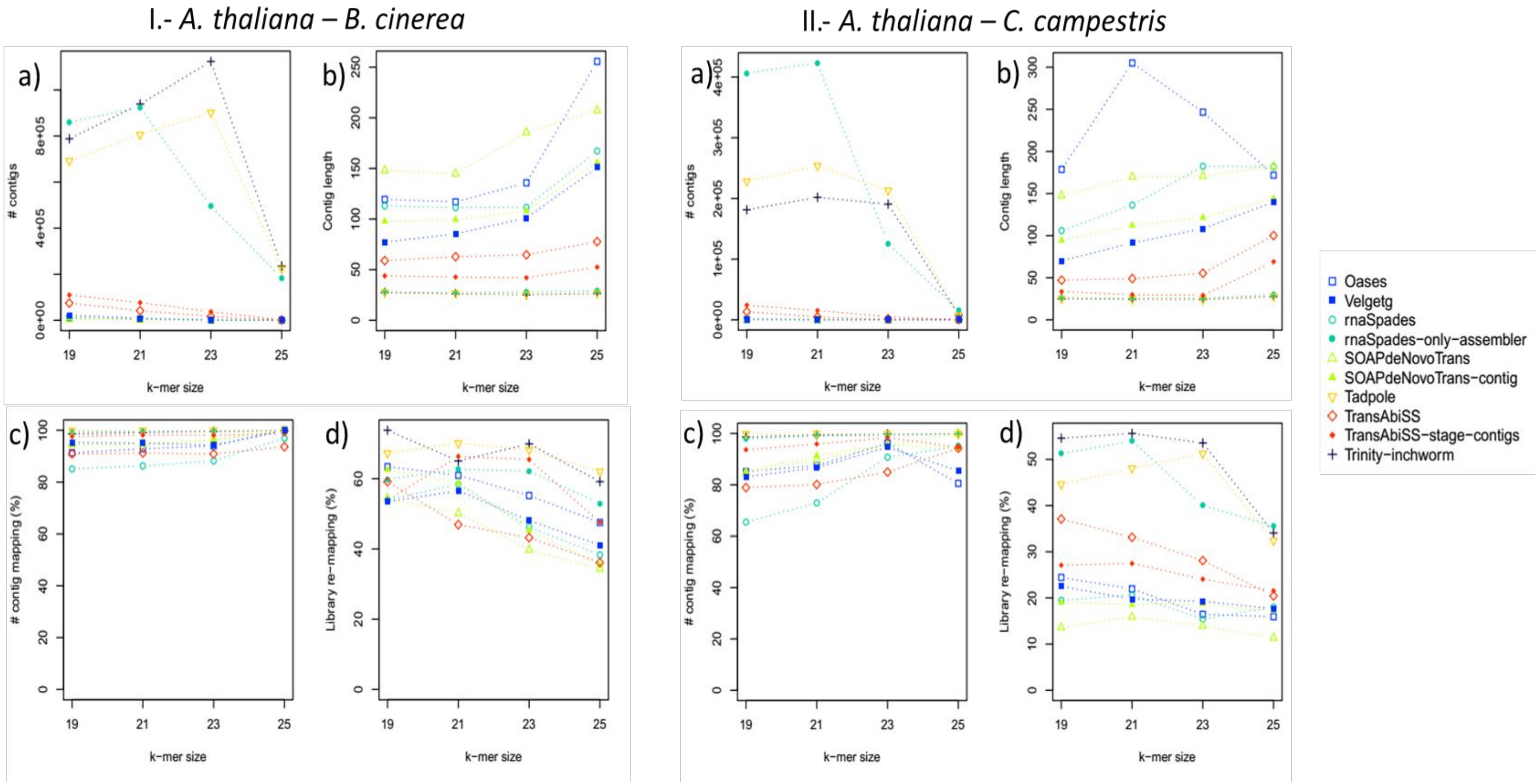


**Supplementary Figure 4. Evaluation of the *de novo* assembly tools at different k-mer sizes.** The effect of using different k-mer sizes during *de novo* assembly is shown for four interaction datasets. a) Total number of generated contigs, b) average length of contigs, c) percent of contigs that map perfectly to the reference genomes, d) percent of initial libraries that map perfectly to the contigs.

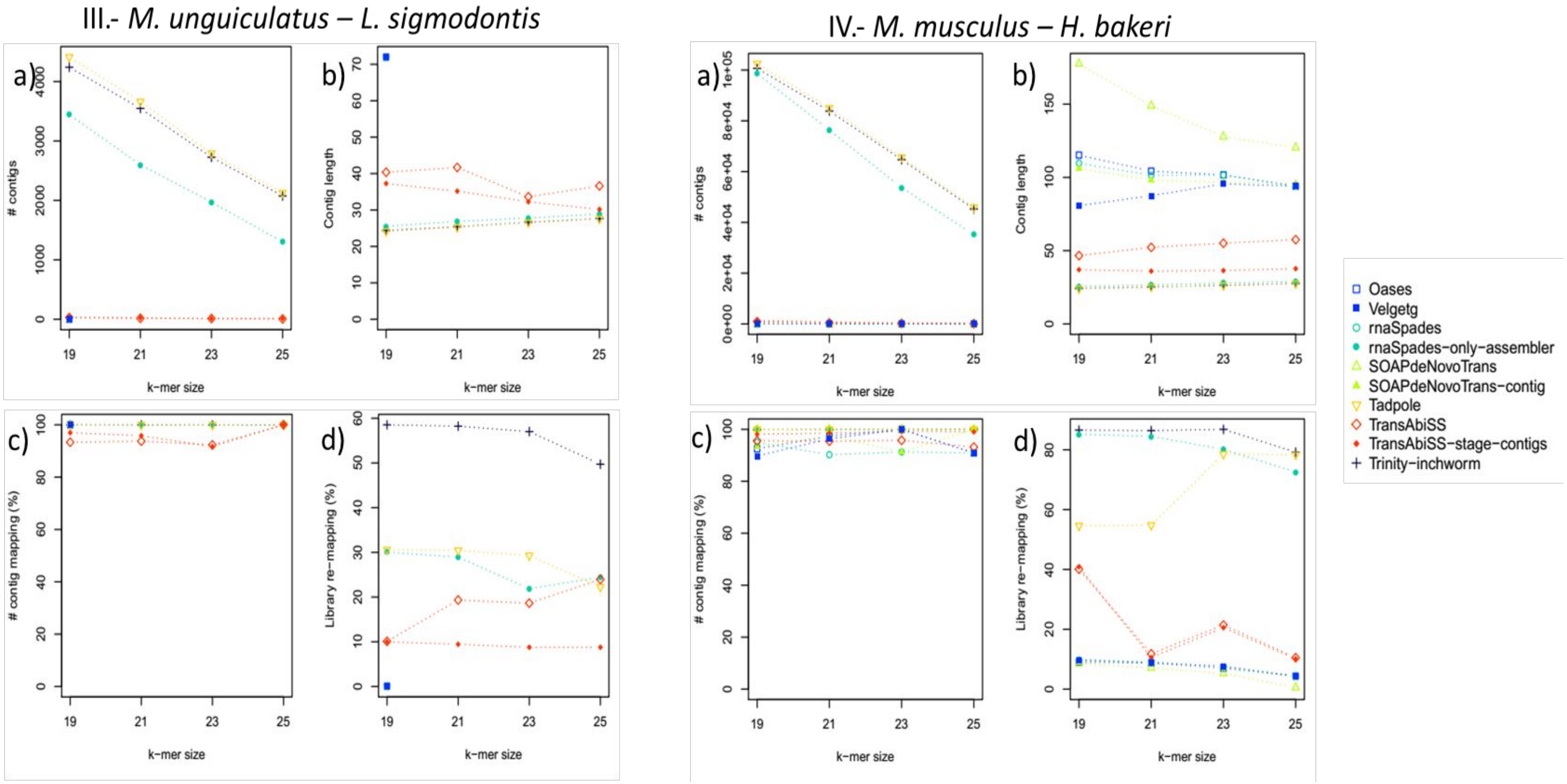


**Supplementary Figure 4. (Continued)**

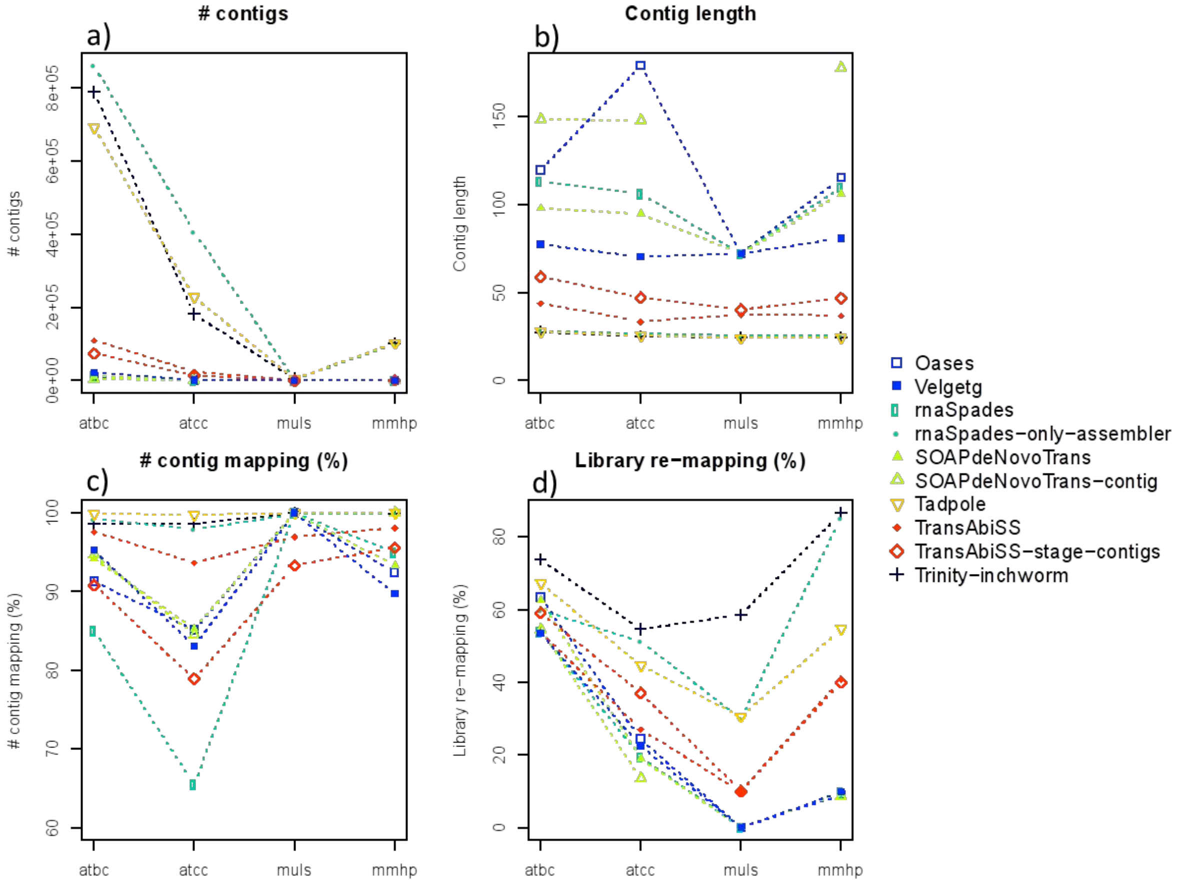


**Supplementary Figure 5. Evaluation of RNA-Seq *de novo* assembly tools using sRNA-Seq data.** The performance of two assembly variants (full transcript reconstruction and "k-mer extension" stage (see Methods)) is shown on our four interaction datasets. a) Total number of generated contigs, b) average length of contigs, c) percent of contigs that map perfectly to the reference genomes, d) percent of initial libraries that map perfectly to the contigs. atbc=A. thaliana and B. cinerea, atcc=A. thaliana and C. campestris, mmhp=M. musculus and H. bakeri, muls=M. unguiculatus and L. sigmodontis.

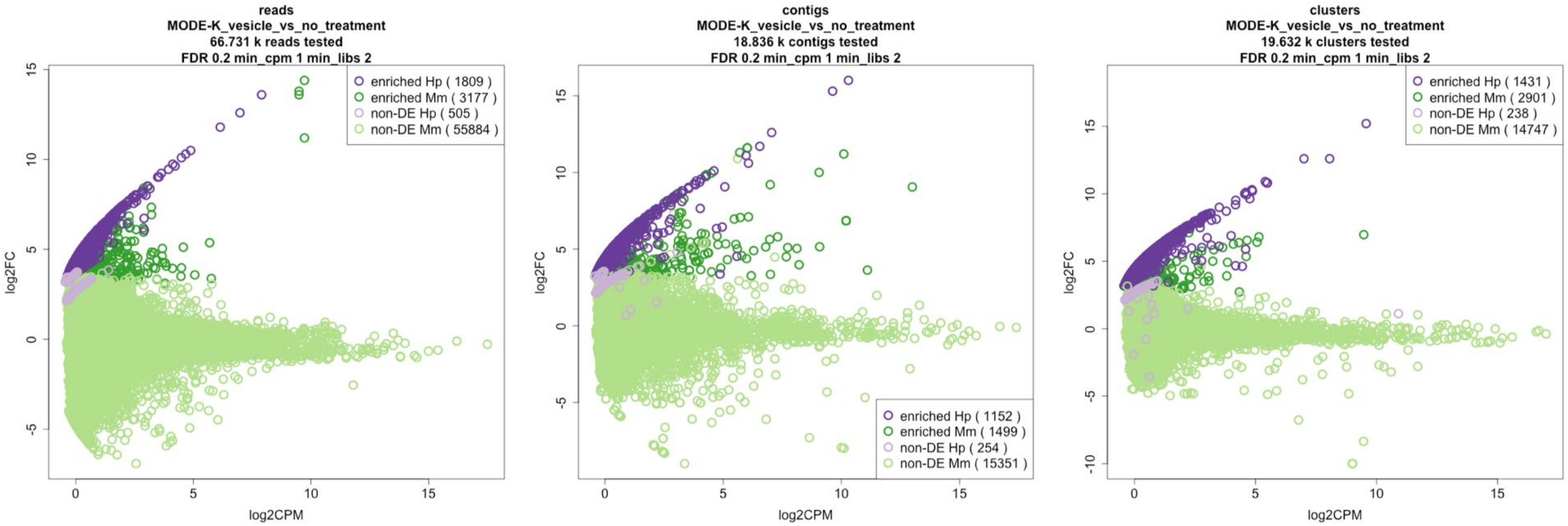


**Supplementary Figure 6. Mean abundance plots of differential expression analysis for all three strategies.** a) Reads or baseline analysis (no assembly) b) *de novo* assembled contigs c) genome-guided assembled clusters. In all cases X-axis represents expression (log2 counts per million, CPM), Y-axis represents fold change EV-treated vs EV-untreated. Differentially expressed (DE) *H. bakeri* sequences are shown in dark purple, non-DE *H. bakeri* sequences are shown in light purple, DE *M. musculus* sequences are show in dark green and non-DE *M. musculus* sequences are shown in light green. The number of sequences in each described set is shown in parenthesis in the legend. In all cases a FDR <= 0.2 was considered as DE threshold. Lowly expressed sequences, less than one CPM in at least two libraries, were filtered prior to differential expression analysis.

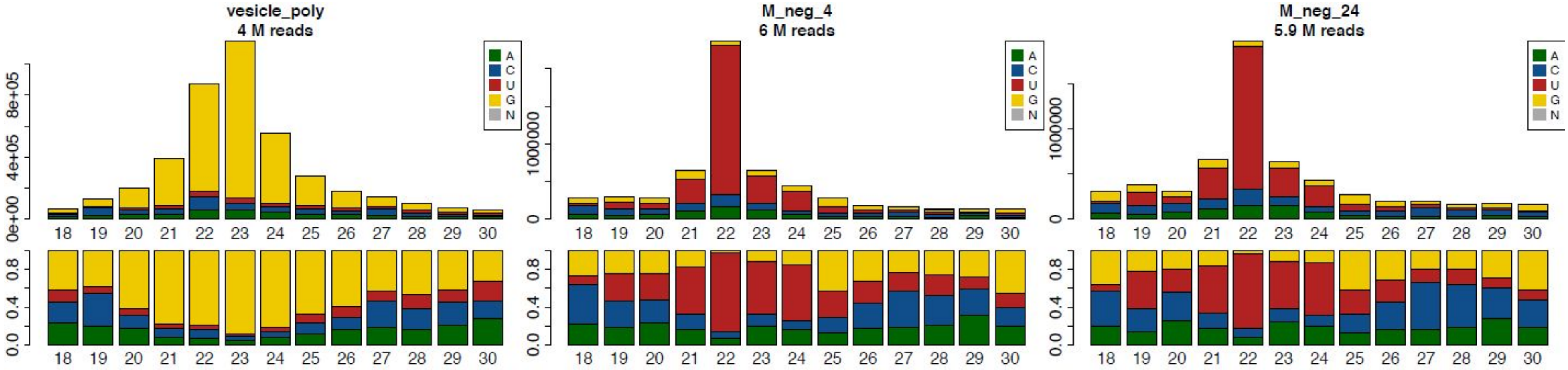


**Supplementary Figure 7. sRNA profile for *H. baker*i vesicles and *M. musculus* MODE-K cells.** First nucleotide and length distribution for polyphosphatase-treated *H. bakeri* vesicles a) and *M. musculus* MODE-K libraries at b) 4 and c) 24 hours incubation without treatment. The average of replicate libraries is shown in this figure.
